# Metabolic flexibility in response to resistance training plus photobiomodulation therapy in sedentary elderly women: a pilot study

**DOI:** 10.64898/2026.05.23.727356

**Authors:** Ronaldo Ferreira Moura, Emanuelle Fernandes Prestes, Danielle Garcia de Araujo, Gislane Ferreira de Melo, Thiago dos Santos Rosa, James W. Navalta, Wilson Max Almeida Monteiro de Moraes, Cleber Ferraresi, Jonato Prestes

**Author notes:** These authors contributed equally to this work (Ronaldo Ferreira Moura, Emanuelle Fernandes Prestes, Danielle Garcia de Araujo, Jonato Prestes). These authors also contributed equally to this work (Gislane Ferreira de Melo, Thiago dos Santos Rosa, James W. Navalta, Wilson Max Almeida Monteiro de Moraes, Cleber Ferraresi).

## Abstract

The study aimed to evaluate the metabolic flexibility of sedentary elderly women in response to resistance training (RT) plus photobiomodulation therapy (PBMT) or RT alone, after two months of intervention. Nineteen elderly women were allocated into two groups, RT (n = 9, 68.44 ± 5.27 years old) and RT plus PBMT (RTPT) (n = 10, 69.40 ± 5.21 years old). The RTPT group received the PBMT, while for the RT group, the equipment was turned off. An incremental treadmill test together with a gas analyzer was performed to record variables such as heart rate (HR), oxygen consumption (VO_2_), carbon dioxide production (VCO_2_) and power output (PO) at the anaerobic threshold (AT), respiratory compensation point (RCP) and maximal oxygen consumption (VO_2max_), and thus indirectly verify metabolic flexibility. In comparison with baseline RTPT displayed significant differences in VO_2max_ (pre: 18.32 ± 3.01; post: 21.89 ± 2.35), carbohydrate oxidation (CHox) (pre: 1.54 ± 0.61; post: 2.45 ± 0.91), CHox/FFM (fat-free mass) (pre: 36.12 ± 12.28; post: 55.92 ± 16.74) and energy expenditure normalized to FFM, EE/FFM (pre: 147.00 ± 49.95; post: 227.56 ± 68.08) during maximum effort incremental testing, while the RT group did not demonstrate a significant difference in these variables.The intervention with RT plus PBMT seems to result in a positive impact on metabolic variables in sedentary elderly women when compared with RT alone, making this approach a viable alternative to improve VO_2max_, CHox, CHox/FFM and EE/FFM during maximal effort testing.

**Author summary:** An aging-related decline in metabolic flexibility may reduce the ability to efficiently use energy during exercise, contributing to lower physical capacity and increased health risks in older adults. Resistance training is widely recommended for elderly populations, but additional strategies may further improve metabolic and physiological adaptations. This study investigated whether combining resistance training with photobiomodulation therapy could improve metabolic responses in sedentary elderly women more effectively than resistance training alone. Participants completed two months of supervised training, and metabolic responses were evaluated during an incremental exercise test. Women who received both resistance training and photobiomodulation therapy showed greater improvements in maximal oxygen consumption, carbohydrate oxidation, and energy expenditure during maximal exercise compared with those who performed resistance training alone. These findings suggest that photobiomodulation therapy may enhance the physiological adaptations associated with resistance training. The results indicate that combining resistance training with photobiomodulation therapy may represent a promising non-invasive strategy to improve exercise metabolism and functional capacity in sedentary elderly women. Further studies with larger samples are needed to confirm these findings and clarify the mechanisms involved.

## Introduction

Metabolic flexibility is a broad term that defines an organism’s ability to switch between energy substrates in response to different conditions and stimuli, such as circadian variation, use of pharmacological compounds, sleeping, fasting, feeding, rest, exercise and environmental fluctuation, aiming to match fuel availability with energy demand [1,2]. Metabolic flexibility takes place across various levels, including substrate, cellular, tissue, organ, and the entire organism. It is influenced by factors such as disease, diet, body composition, and a range of other (epigenetic) factors [2].

Previous studies have demonstrated that young and physically active people exhibit better metabolic flexibility [3,4] compared to those who are older and sedentary [4]. Conversely, metabolic inflexibility is associated with certain pathological conditions common with advancing age such as obesity, metabolic syndrome, type 2 diabetes, cardiovascular diseases, cancer, and sarcopenia [2,5].

Considering metabolic flexibility as a contributing factor to aging and associated comorbidities, its evaluation could offer valuable insights into delaying the onset of age-related diseases and extending the health span [2]. Moreover, metabolic inflexibility leads to mitochondrial dysfunction and cellular alterations that favor a glycolytic phenotype. This condition is strongly associated with aging and contributes to the release of pro-inflammatory cytokines, proteases, and growth factors, which exert significant local and systemic effects, such as inflammation and metastasis [6,7].

The mechanisms underlying metabolic flexibility in older adults remain poorly understood, as research in late middle-aged and elderly populations is limited, while sedentary lifestyle is a major driver of age-related metabolic dysfunction, which becomes more pronounced in older individuals [2]. In this sense, exercise serves as a preventive strategy to improve metabolic flexibility and promote healthy aging [8].

However, studies that have linked metabolic flexibility to chronic exercise interventions are extremely rare. Most studies to date measure acute responses with physical testing protocols carried out with young and middle-aged participants [9,10] and typically people who identify as male [9,11,12]. A study by Ortmeyer et al. [13] examined the effects of 6 months of weight loss versus aerobic exercise training plus weight loss on fat and skeletal muscle markers of fatty acid metabolism in postmenopausal overweight/obese women, normal and impaired glucose tolerant. An improvement in skeletal muscle fatty acid metabolism was reported only in the aerobic exercise plus weight loss group, suggesting that in the elderly, metabolic aspects can still be trained. This highlights the potential of exercise as an intervention to improve metabolic flexibility. Both resistance (RT) and endurance training can provide considerable metabolic health benefits, such as increased mitochondrial content and improvements in glycemic control [14].

To our knowledge, few studies have investigated the chronic effects of RT on metabolic flexibility [15] and no one investigated the effects of RT, especially associated with other tools such as photobiomodulation therapy (PBMT). In this context, PBMT has shown to be an interesting resource that can be applied to skeletal muscle tissue immediately before or after an exercise session to enhance muscle strength or improve resistance to fatigue [16-18]. The PBMT acts on biological tissues through the absorption of light by chromophores in cells, with cytochrome c oxidase (Cox) being the principal and most studied mitochondrial light-absorbing protein involved in the process of photobiomodulation [19,20]. The light used in PBMT can be emitted by diode lasers or LEDs (light-emitting diodes) with low-power (mW) and low-intensity (mW/cm^2^) to avoid tissue heating or any kind of tissue damage [21]. In this context, the light emitted by these light sources present very similar effects when the light parameters used are similar [22]. Furthermore, the fact that cells with many mitochondria and high metabolic activity such as skeletal muscle tissue are highly responsive to light [16,23], makes PBMT an interesting and non-invasive therapy to be investigated in the context of metabolic flexibility.

Scientific literature revealed that PBMT can increase mitochondrial activity, resulting in greater ATP (Adenosine Triphosphate) production and consequently better utilization of energy substrates [23,24], and that greater mitochondrial volume density and better mitochondrial quality can activate Adenosine Monophosphate-activated Protein Kinase (AMPK) in the long term [14,25].The acute activation of AMPK reduces glycogen and protein synthesis and simultaneously facilitates glucose transport and fatty acid oxidation [14,26]. Furthermore, AMPK can also be considered an important regulator of exercise-induced effects on metabolic flexibility [27,28]. The justification for the use of PBMT combined with RT was the search for more pronounced training responses on metabolic flexibility, based on the description in the previous paragraph.

Based on these observations and considering that metabolic flexibility is extremely important to maintain energy homeostasis [2], the attention given in recent years to the metabolic influence on aging and lifespan, as well as the impact that physical exercise has on metabolic flexibility and aging, the aim of the present study was to evaluate the metabolic flexibility of sedentary elderly women in response to RT plus PBMT and RT alone, after two months of intervention. The initial hypothesis was that both RT, and RT plus PBMT, would show positive responses regarding metabolic flexibility, but that TF combined with TFBM would show more pronounced responses in metabolic flexibility than TF alone.

## Materials and Methods

### Subjects and ethical approval

The study began with 22 elderly women participating in other university projects recruited by invitation. So, the participants were randomly assigned to two groups of 11 participants each. However, three participants dropped out during the intervention: one due to a family member’s illness, another due to the death of a sister, and the third because she lived far away and was unable to arrive at the scheduled training time, being two in resistance training (RT) group and one in the resistance training and photobiomodulation therapy (RTPT) group. At the end, nineteen participants remained: nine and ten in the RT and RTPT group, respectively (Table 1). After answering a questionnaire about their health status and level of physical activity, all participants were considered sedentary [29]. The inclusion criteria were: a) 60 years of age or older at the time of the screening; b) no type of regular physical training; c) no contraindication to practicing physical exercise and d) no musculoskeletal injuries. The exclusion criteria were: a) history of metabolic/endocrine (diabetes), or renal disease, myocardial infarction, obesity or respiratory or neuromuscular disease verified through an anamnesis; b) recent surgery and c) the use of hormone replacement therapy. Participants were informed of the risks and benefits of the study before participating, and provided written informed consent before inclusion. Most participants reported regular use of antihypertensive medication; some reported using pharmacological treatment for diabetes, and three reported using medication for dyslipidemia. Despite the presence of these chronic conditions, the overall health status of the elderly women evaluated was considered good for their age group. This study was approved by the Catholic University of Brasilia Institutional Review Board (protocol number: 75254823.4.0000.0029), and was conducted in accordance with the Declaration of Helsinki [30].

**Table 1.** Descriptive characteristics the participants.

| | RT | | RTPT | | Group | P and effect size ($\omega^2$) | | | | |
| --- | --- | --- | --- | --- | --- | --- | --- | --- | --- | --- |
| | Pre | Post | Pre | Post | | $\omega^2$ | Time | $\omega^2$ | Group* Time | $\omega^2$ |
| Age | 68.44 ± 5.27 | 68.67 ± 5.43 | 69.40 ± 5.21 | 69.90 ± 5.04 | 0.52 | 0.00 | 0.83 | 0.00 | 0.94 | 0.00 |
| Height | 153.00 ± 4.58 | 152.93 ± 4.92 | 155.45 ± 6.21 | 155.17 ± 5.98 | 0.20 | 0.00 | 0.92 | 0.00 | 0.95 | 0.00 |
| Weight | 61.91 ± 11.61 | 59.53 ± 9.48 | 64.57 ± 13.63 | 64.64 ± 13.77 | 0.34 | 0.00 | 0.78 | 0.00 | 0.76 | 0.00 |
| BMI | 26.44 ± 4.68 | 25.57 ± 4.70 | 26.69 ± 5.16 | 26.79 ± 5.16 | 0.65 | 0.00 | 0.81 | 0.00 | 0.76 | 0.00 |
| FFM (kg) | 39.69 ± 4.80 | 40.84 ± 4.77 | 42.34 ± 4.56 | 43.41 ± 4.61 | 0.09 | 0.05 | 0.47 | 0.00 | 0.98 | 0.00 |
| Fat % | 35.91 ± 6.90 | 36.12 ± 7.23 | 36.03 ± 8.11 | 36.50 ± 8.03 | 0.92 | 0.00 | 0.89 | 0.00 | 0.96 | 0.00 |
| $VO_{2max}$ (ml/min) | 18.98 ± 5.03 | 21.16 ± 4.88 | 18.32 ± 3.01 | 21.89 ± 2.35 | 0.69 | 0.00 | 0.96 | 0.00 | 0.46 | 0.00 |

Data are shown as average ± SD. RT: resistance training, RTPT: resistance training plus photobiomodulation therapy, BMI: body mass index, FFM: fat-free mass, $VO_{2max}$ : maximal oxygen uptake. $\omega^2$ : omega square estimate of effect size.

### Study design

The study is begun with the first visit to the laboratory for informed consent, assessment of body composition through whole-body Dual-energy X-ray Absorptiometry (DXA), and completion of the 10-repetition maximum (RM) test. 48 hours later, the maximum incremental test was performed on a treadmill. The following week, corresponding to the first week of intervention, consisted of adaptation and learning sessions for resistance training exercises. Then, from the second week onwards, the training protocol was initiated, in which the participants were allocated to one of the following groups: resistance training (RT) or resistance training plus photobiomodulation therapy (RTPT). The data were compared to ensure that the groups receiving the intervention had no significant differences between the variables assessed. Thus, as participants were counterbalanced in the groups based on initial strength values and PBMT was applied to both groups for 5 minutes and 22 seconds on the quadriceps femoris of both thighs, with the difference that the RTPT group effectively received the PBMT protocol, whereas for the RT group, PBMT was used with the equipment turned off, characterizing the study as a sham. The study was conducted blindly and included only the evaluators. The intervention protocol lasted for a duration of two months (24 sessions). There was data pairing so that the groups received the intervention without significant differences between the variables evaluated. Finally, after the end of the intervention period, the last visit consisted assessment of body composition and the incremental treadmill test to verify changes in body composition and metabolic data (Figure 1)

**Figure 1.**
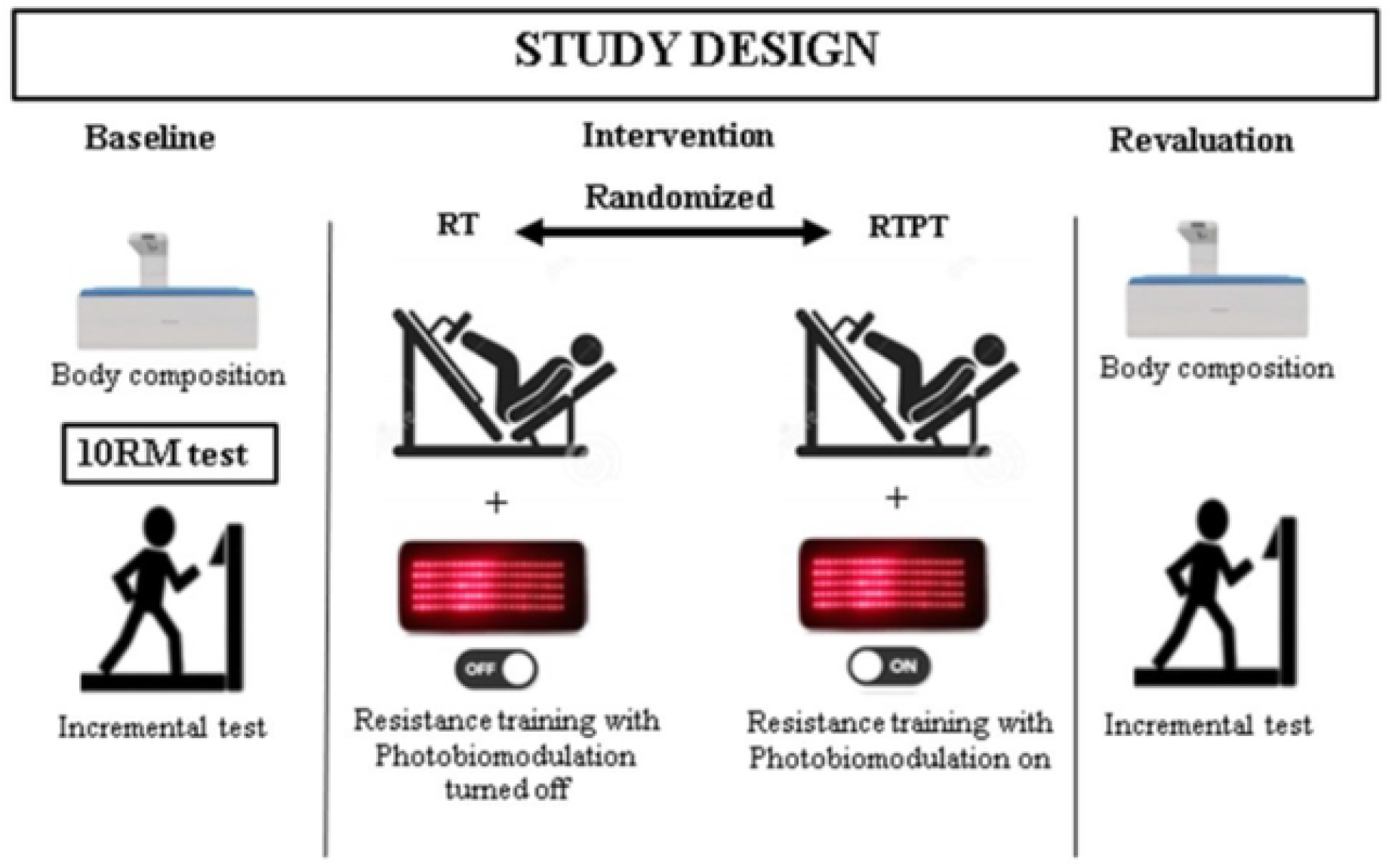
Overview of the study design. Note = RT: resistance training, RTPT: resistance training plus photobiomodulation

### Body composition assessment

Body mass was measured with the volunteer barefoot, wearing light clothing and using a digital scale (Welmy-W110H, Sao Paulo, Brazil) with a capacity of 150 kg and a sensitivity of 100 g. Height was measured using a wall stadiometer (Sanny, Sao Paulo, Brazil), with a capacity of 2,200 mm and a sensitivity of 1 mm.

Body composition analysis was carried out using the DXA technique (Lunar Corp. Madson, 219 WI USA, DPX-IQ 2446). Body mass was assessed, as were fat mass, fat-free mass (FFM) and percentage of fat mass (%Fat). All participants performed the test wearing light sports clothing. During the test, participants placed their body and hands in a supine position, and their feet was secured by a strap. Participants were instructed not to move or talk while the scanner performed measurements. The tests included a complete scan of volunteers’ body for approximately 17 minutes, with the device always adjusted and operated by a technically trained professional. The scan was calibrated every two days using the phantom supplied by the manufacturer.

### Nutritional assessment

Although there was no nutritional assessment or intervention, a self-selected diet was carried out. The participants were instructed to maintain their normal eating habits and on the day of the tests: could eat 2 hours before, no smoking 2 hours before; abstain from alcohol, coffee, tea, chocolates and soft drinks at least 8 hours before; and were instructed not to perform physical exercises on test day.

### Incremental exercise test

The incremental treadmill test was performed twice, before and after two months of intervention. To assess cardiopulmonary condition, a gas analyzer (Cortex Metalyzer 3B, Leipzig, Germany) and a mask (Hans Rudolph) were used for the direct measurement of oxygen uptake (VO_2_) and ventilatory parameters. Additionally, the incremental test was conducted on a treadmill (IMBRAMED, Super ATL-32km/h, Porto Alegre, Brazil), with an estimated duration of 10 minutes (Ramp protocol). During testing with all participants, the ambient temperature was maintained between ∼24 and 25 °C.

Before the test began, all procedures were clearly explained to the participants. The test began with an initial speed of 3 km/h and an incline of 3%, with a final expected speed of 6 km/h and an incline of 12%. After the start, speed and incline increased gradually: 0.1 km/h every 10 seconds and 0.5% every 40 seconds, respectively. During the test, volunteers were continuously encouraged through verbal motivation, and both inability to continue the activity and voluntary withdrawal were observed. The test was interrupted in case the individual requested it for fatigue or presented any of the absolute criteria for the test interruption, according to the guidelines of the American Heart Association [31].

To ensure data accuracy, variables such as heart rate (HR), oxygen uptake (VO_2_), carbon dioxide production (VCO_2_), and power output (PO) were recorded in three distinct phases of progressive cardiorespiratory effort: anaerobic threshold (AT), respiratory compensation point (RCP) and maximal oxygen uptake (VO_2max_). For better understanding, LA, CRP and VO_2max_ were classified into three stages: S1, S2 and S3, respectively (Table 2). These parameters were analyzed to identify VO_2_, CO_2_, and non-buffered acidosis at different effort levels [32]. AT was determined based on the second ventilatory threshold, which coincide with the non-linear increase in ventilation and CO_2_ production [33], while RCP was determined by the sudden increase in final O_2_ pressure without a reduction in final CO_2_ pressure, reflecting respiratory alkalosis [34]. To monitor HR beat by beat during the test, an electrocardiographic recording (ECG Elite Micromed, Brasilia, Brazil) was used. All recordings were evaluated by a cardiologist to detect potential cardiovascular abnormalities that could prevent test execution. The test was considered maximal if two of the following three criteria were met: HR ≥ 95% of the theoretical maximum heart rate (220 - age), respiratory exchange ratio ≥ 1.10 and VO_2_ stabilization despite increasing exercise intensity [35]. The highest VO_2_ value achieved during the test was considered as VO_2max._

**Table 2.** Values of metabolic markers in the incremental exercise test in pre and post intervention.

| | RT | | RTPT | | P and effect size ($\omega^2$) | | | | | |
| --- | --- | --- | --- | --- | --- | --- | --- | --- | --- | --- |
| | Pre | Post | Pre | Post | Group | $\omega^2$ | Time | $\omega^2$ | Group*<br>Time | $\omega^2$ |
| **S1 - Anaerobic threshold** |  |  |  |  |  |  |  |  |  |  |
| HR (bpm) | 104.44 ± 19.70 | 108.67 ± 15.67 | 106.60 ± 11.92 | 108.10 ± 9.29 | 0.87 | 0.00 | 0.55 | 0.00 | 0.77 | 0.00 |
| $VO_2$ (ml/min/kg) | 8.40 ± 3.79 | 11.11 ± 4.10 | 11.82 ± 3.25 | 12.95 ± 2.96 | 0.03† | 0.10 | 0.10 | 0.04 | 0.50 | 0.00 |
| FATox (g/min) | 0.19 ± 0.11 | 0.20 ± 0.08 | 0.28 ± 0.11 | 0.30 ± 0.12 | 0.009† | 0.15 | 0.66 | 0.00 | 0.90 | 0.00 |
| CHox (g/min) | 0.17 ± 0.08 | 0.34 ± 0.20 | 0.25 ± 0.18 | 0.30 ± 0.24 | 0.79 | 0.00 | 0.08 | 0.06 | 0.30 | 0.00 |
| PO (W) | 48.06 ± 15.00 | 54.98 ± 17.28 | 58.70 ± 17.28 | 67.76 ± 20.85 | 0.051 | 0.08 | 0.18 | 0.02 | 0.85 | 0.00 |
| **S2 - Respiratory compensation point** |  |  |  |  |  |  |  |  |  |  |
| HR (bpm) | 129.67 ± 21.91 | 130.44 ± 15.91 | 127.30 ± 13.38 | 136.30 ± 14.06 | 0.75 | 0.00 | 0.37 | 0.00 | 0.45 | 0.00 |
| $VO_2$ (ml/min/kg) | 15.64 ± 4.78 | 18.50 ± 5.11 | 16.38 ± 2.17 | 18.76 ± 2.60 | 0.69 | 0.00 | 0.04‡ | 0.09 | 0.85 | 0.00 |
| FATox (g/min) | 0.16 ± 0.12 | 0.09 ± 0.05 | 0.11 ± 0.08 | 0.10 ± 0.07 | 0.47 | 0.00 | 0.09 | 0.05 | 0.30 | 0.00 |
| CHox (g/min) | 0.84 ± 0.47 | 1.23 ± 0.44 | 1.11 ± 0.37 | 1.37 ± 0.36 | 0.13 | 0.03 | 0.02‡ | 0.11 | 0.65 | 0.00 |
| PO (W) | 98.32 ± 36.24 | 125.46 ± 38.17 | 119.36 ± 47.87 | 144.67 ± 41.84 | 0.14 | 0.03 | 0.06 | 0.07 | 0.95 | 0.00 |
| **S3 - Maximal oxygen uptake** |  |  |  |  |  |  |  |  |  |  |
| HR (bpm) | 143.44 ± 22.48 | 153.11 ± 22.32 | 141.10 ± 15.57 | 152.80 ± 15.13 | 0.83 | 0.00 | 0.09 | 0.05 | 0.87 | 0.00 |
| $VO_2$ (ml/min/kg) | 18.98 ± 5.03 | 21.16 ± 4.88 | 18.32 ± 3.01 | 21.89 ± 2.35* | 0.98 | 0.00 | 0.03* | 0.10 | 0.59 | 0.00 |
| FATox (g/min) | 0.02 ± 0.16 | -0.11 ± 0.11 | 0.005 ± 0.13 | -0.22 ± 0.26* | 0.30 | 0.00 | 0.004* | 0.18 | 0.41 | 0.00 |
| CHox (g/min) | 1.42 ± 0.69 | 1.91 ± 0.52 | 1.54 ± 0.61 | 2.45 ± 0.91* | 0.15 | 0.02 | 0.004* | 0.18 | 0.37 | 0.00 |
| PO (W) | 135.66 ± 38.54 | 170.46 ± 45.49 | 151.47 ± 59.90 | 193.30 ± 46.90 | 0.23 | 0.01 | 0.021‡ | 0.12 | 0.83 | 0.00 |

Data are shown as average ± SD. RT: resistance training, S: Stage, RTPT: resistance training plus photobiomodulation therapy, HR: heart rate, $VO_2$ : oxygen uptake. FATox: fat oxidation, CHox: carbohydrate oxidation. PO: power output, $\omega^2$ : omega square estimate of effect size. †: significant difference between groups ( $p \leq 0.05$ ), \*: significant differences between time points ( $p \leq 0.05$ ), ‡: statistical difference became null after pairwise comparisons with Bonferroni correction.

### Resistance training protocol

For purposes of training adaptation, all subjects completed familiarization training for one week, over two sessions, allowing for the equalization of groups and counterbalancing subjects according to muscular strength, which was verified through repetition maximum testing (10RM), as described below. After the adaptation period, a linear periodization protocol was used. The full body method using the alternating segment approach was chosen for the training protocol. Priority was given to multi-joint exercises of pushing and pulling patterns for the lower body and upper body. The exercises were: horizontal leg press, dumbbell bench press, leg extension, seated leg curl, seated cable row and seated plantar flexion.

The training lasted two months, with three weekly sessions (24 sessions). The average duration of each session was 50 minutes. The duration of each repetition was 3-4s, including concentric and eccentric muscle actions. All sessions took place in the afternoon and were supervised by a professional with experience in RT.

The choice of load percentage was based on a previous study that demonstrated improvements in mitochondrial metabolism and metabolic flexibility [36]. During the linear periodization protocol, in the initial 2 weeks, 3 sets of 10 repetitions were performed at 70% of 10RM; from the 3rd and 4th week, 3 sets of 10 repetitions were performed at 80% of 10RM; in the 5th and 6th week, 3 sets of 10 repetitions were performed at 90% of 10RM; and in the 7th and 8th weeks, workout consisted of 3 sets of 10 repetitions at 100% of 10RM. In all phases, the interval between sets and exercises was 60 seconds.

### 10 RM protocol

Subjects began the protocol by performing a warm-up set consisting of 5 to 10 repetitions, using between 40% and 60% of the estimated maximum load for 10RM. After a 5-minute rest, they completed the 10RM test. At the beginning of the test, the maximum load that each participant believed they could lift for 10 repetitions was used. If the load was not adequate on the first attempt, the researcher adjusted the weight and then the participant performed a new test after 2-minutes of rest. After the second attempt, if the load was still not correct, a minimum rest of 5 minutes was established before performing the last attempt.

Each participant was allowed a maximum of three attempts per load variation. All participants were able to determine their maximum load for 10 RM on the first day of testing. After determining the loads, subjects were instructed to rest for 48 hours before any additional training or assessment sessions (10RM tests were repeated 48-72 h later). This was used to calculate reproducibility of the tests, with an overall R = 0.97-0.99. The 10 RM tests were performed for all exercises carried out in the workouts, with a 2-minute rest interval between each exercise.

### Photobiomodulation therapy (PBMT)

For the PBMT intervention, a phototherapy device composed of low-intensity LEDs using four flexible “blankets” containing 264 individual commercial LED light emitters (BODY 264 PAD RED Sonoita, United States) each. Each “blanket” featured 120 LEDs (5 rows of 24 LEDs) that emitted light in the red range (635 ± 10 nm) and 144 LEDs (6 rows of 24 LEDs) emitting in the infrared range (880 ± 20 nm) with an irradiation surface area of 333.5 cm^2^ (11.5 × 29 cm). Each red LED had 1.2 mW optical power, and each infrared LED had 15 mW both with 4.7 kHz pulse frequency measured with an optical and energy meter (PM100D Thorlabs®, Newton, United States) equipped with an S130C sensor. The total optical power (W) of each “blanket” was 2,304 W [37].

The “blankets” were positioned perpendicularly (90°) to the skin of the quadriceps femoris region. The PBMT was applied bilaterally to the thighs (rectus femoris, vastus lateralis and vastus medialis) in contact mode with light pressure, and two “blankets” on each thigh, while the volunteers were lying in the supine position. The time of radiation (PBMT application) per region was 322 s (5’22’’), delivering ∼741 J (∼2.22 J/cm^2^) per “blanket”, surpassing the minimal local dose of 1 J/cm^2^ for exercise performance enhancement and postexercise recovery suggested previously [38]. Each red LED emitted ∼0.38 J, while infrared LED emitted ∼4.83 J, totaling 1.080 J applied on each thigh 10 minutes after each exercise session, similarly to previous studies [17,39]. PBMT was performed three times a week, 10 minutes after the resistance training session, always in the afternoon, between 3 and 5 pm. The same procedure was carried out with both groups (RT and RTPT), with RT group receiving sham therapy (the PBMT device was turned off). PBMT was applied by the same researcher in all sessions.

The potential risks of PBMT use are limited to a few generally mild side effects, which may include skin irritation, itching, and redness. These are not very harmful and do not increase the temperature of the target tissue [40]. To minimize these effects, we asked participants to report any of these symptoms, and if any were detected, the trial would be immediately stopped. Fortunately, no such reactions occurred during the trials.

### Measurements

Heart rate (HR) was measured beat by beat and registered every 10 seconds throughout the test by electrocardiographic recording (ECG Elite Micromed, Brasilia, Brazil) and maximum heart rate was considered as the highest heart rate value obtained during the test.

The volume and composition of expired gas were analyzed breath by breath using a gas analyzer (Metalyzer 3B, CPX System, Cortex, Germany), previously calibrated according to the manufacturer instructions. oxygen uptake (VO_2_) and excretion of carbon dioxide (VCO_2_) were calculated every 10 seconds from the gas analyzer software. VO_2max_ was established as the highest value achieved during the incremental exercise test. Fat oxidation (FATox) and carbohydrate oxidation (CHox) were calculated from the respiratory quotients (RQ) using standard indirect calorimetry equations [41]. Both FATox and CHox were used to indirectly assess metabolic flexibility.

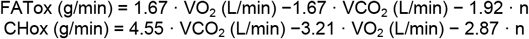

Where n represents negligible urinary nitrogen excretion. In the above calculations, CHO and FAT oxidation are based on a VO_2_ (STPD) of 2.500 L·min^-1^ and a VCO_2_ (STPD) of 2.250 L·min^-1^ and negligible protein oxidation (n = 0) [42].

For the analysis, FATox and CHox rates were normalized to kilogram of FFM. Maximum fat oxidation (MFO) was determined as the highest FATox value during the test. MFO/FFM was determined by normalizing MFO to FFM. In addition, variables such as: VO_2_, percentage of VO_2max_ (%VO_2max_), HR, percentage of HRmax (%HR_max_), power output (PO), and PO normalized to FFM (PO/FFM) were determined at the intensity corresponding to MFO. Energy expenditure (EE) was normalized to FFM (EE/FFM) and determined from FATox/FFM and CHox/FFM, using the predictive equation below [42].

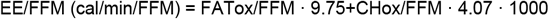

### Statistical analysis

A two-way ANOVA was conducted to examine the effects of group and time on the dependent variables. Data are presented as mean ± standard deviation unless otherwise stated. Residual analysis was performed to test the assumptions of the two-way ANOVA. Outliers were assessed by inspecting a boxplot, normality was evaluated using Shapiro-Wilk’s test for each cell of the design, and homogeneity of variances was assessed using Levene’s test. No outliers were detected, residuals were normally distributed (*p* >.05), and homogeneity of variances was confirmed (*p* >.05).

When a statistically significant interaction between group and time on the dependent variables was observed, an analysis of simple main effects for group and time was conducted, with statistical significance adjusted using the Bonferroni correction. For effect size estimation, omega squared (ω^2^) was used, as it is less biased than eta squared (η^2^), particularly in small samples. Values of < 0.04, 0.04–0.24, 0.25–0.63, and ≥ 0.64 were interpreted as trivial, small, medium, and large, respectively [43].

Based on a previous study, the variables used to calculate power were FATox/FFM and Chox/FFM at stage three [3]. To determine the main effect of group, the *a priori* F-test option (ANOVA: Fixed effects, special, main effects, and interactions) was used. For FATox/FFM, the effect size was 0.15, with an alpha error probability of 0.05, power of 0.80, numerator degrees of freedom (df) of 1 (since the group factor had 2 levels), and a total of 4 groups in the design.

For all analyses, the following software programs were used: Jeffreys’s Amazing Statistics Program (JASP 0.19.3, Amsterdam, Netherlands), Statistical Package for the Social Sciences (SPSS 21.0, Chicago, EUA) and G*Power v. 3.1.9.2 (Stuttgart, Germany). For all comparisons, a significance level of α ≤0.05 was adopted [44-46].

## Results

### Descriptive characteristics

No interaction between group*time on age (*F*(1, 34) = 0.007, *p* = 0.94, height *F*(1, 34) = 0.004, *p* = 0.95), body weight (*F*(1, 34) = 0.093, *p* = 0.76, BMI *F*(1, 34) = 0.092, *p* = 0.76), FFM (*F*(1, 34) = 0.001, *p* = 0.98, FAT% *F*(1, 34) = 0.003, *p* = 0.96), and VO_2_max (*F*(1, 34) = 0.56, *p* = 0.46) were observed (Table 1).

### Anaerobic threshold

No interaction between group*time on heart rate (*F*(1, 34) = 0.084, *p* = 0.77), FATox (*F*(1, 34) = 0.016, *p* = 0.90), CHox (*F*(1, 34) = 1.11, *p* = 0.30), VO_2_ (ml/kg/min) (*F*(1, 34) = 0.043, *p* = 0.50), and power output (*F*(1, 34) = 0.034, *p* = 0.85) were observed. However, a main effect of group was verified for VO_2_ (ml/kg/min) (*F*(1, 34) = 5.23, *p* = 0.03), and FATox (*F*(1, 34) = 7.61, *p* = 0.009) (Table 2).

The RTPT displayed a statistically superior mean difference at baseline of 0.10 g/min FATox compared to RT (*p* = 0.049, and small effect size). Also, at baseline the RTPT displayed a statistically superior mean difference of 3.41 VO_2_ (ml/kg/min) than RT (*p* = 0.043, and small effect size) (Table 2).

### Respiratory compensation point

No interaction between group*time on heart rate (*F*(1, 34) = 0.588, *p* = 0.45), FATox (*F*(1, 34) = 1.09, *p* = 0.30), CHox (*F*(1, 34) = 0.21, *p* = 0.65), VO_2_ (ml/kg/min) (*F*(1, 34) = 0.038, *p* = 0.85), and power output (*F*(1, 34) = 0.005, *p* = 0.95) were observed. However, a main effect of time was verified for VO_2_ (ml/kg/min) (*F*(1, 34) = 4.46, *p* = 0.04), and CHox (*F*(1, 34) = 5.84, *p* = 0.02). After pairwise comparisons, with Bonferroni corrections, differences became null for VO_2_ (ml/kg/min) (*p* = 0.12, and p = 0.17 for RT and RTPT, respectively) but the effect size was trivial. Also, after pairwise comparisons, with Bonferroni corrections, differences became null for CHox (*p* = 0.055, for RT) but the effect size was small (Table 2).

### Maximal oxygen uptake

No interaction between group*time on heart rate (*F*(1, 34) = 0.027, *p* = 0.87), FATox (*F*(1, 34) = 0.68, *p* = 0.41), CHox (*F*(1, 34) = 0.83, *p* = 0.37), VO_2_ (ml/kg/min)^-^ (*F*(1, 34) = 0.299, *p* = 0.59), and power output (*F*(1, 34) = 0.050, *p* = 0.83) were observed. However, a main effect of time was verified for VO_2_ (ml/kg/min) (*F*(1, 34) = 5.07, *p* = 0.031), FATox (*F*(1, 34) = 9.19, *p* = 0.004), CHox (*F*(1, 34) = 9.33, *p* = 0.004), and PO (*F*(1, 34) = 5.89, *p* = 0.021) (Table 2).

The RTPT displayed a statistically superior mean difference of 3.57 VO_2_ (ml/kg/min) at the post time point (*p* = 0.050, and small effect size) compared to baseline, and a statistically inferior mean difference of FATox at the post time point (*p* = 0.049, and small effect size) compared to baseline. Also, at the post time point the RTPT displayed a statistically superior mean difference of 0.90 g/min of CHox (*p* = 0.007, and small effect size) compared to baseline. For power output, after pairwise comparison with Bonferroni correction, differences became null (*p* = 0.063) for RTPT, but with a small effect size (Table 2).

### Maximal fat oxidation

No interaction between group*time on MFO (*F*(1, 34) = 0.302, *p* = 0.59), MFO/FFM (*F*(1, 34) = 0.41, *p* = 0.52), VO_2_ (ml/kg/min) (*F*(1, 34) = 0.36, *p* = 0.55), VO_2_/FFM (*F*(1, 34) = 0.84, *p* = 0.36), %VO_2max_ (*F*(1, 34) = 0.067, *p* = 0.80), heart rate (*F*(1, 34) = 0.24, *p* = 0.63), %HR _max_ (*F*(1, 34) = 0.36, *p* = 0.55), PO (*F*(1, 34) = 2.22, *p* = 0.14), and PO/FFM (*F*(1, 34) = 3.02, *p* = 0.09) were observed. However, a main effect of time was verified for %VO_2max_ (*F*(1, 34) = 6.60, *p* = 0.01), and %HR _max_ (*F*(1, 34) = 9.19, *p* = < .001). The RTPT displayed a statistically inferior mean difference of -9.34 %VO_2max_ (*p* = 0.49, and small effect size) at the post time point compared to baseline. Also, RT (*p* = 0.003, and medium effect size) and RTPT (*p* = 0.017, and medium effect size) displayed a statistically inferior mean difference of -9.82 %HR _max_ and -7.26 %HR _max_, respectively at the post time point compared to baseline (Table 3).

**Table 3.**
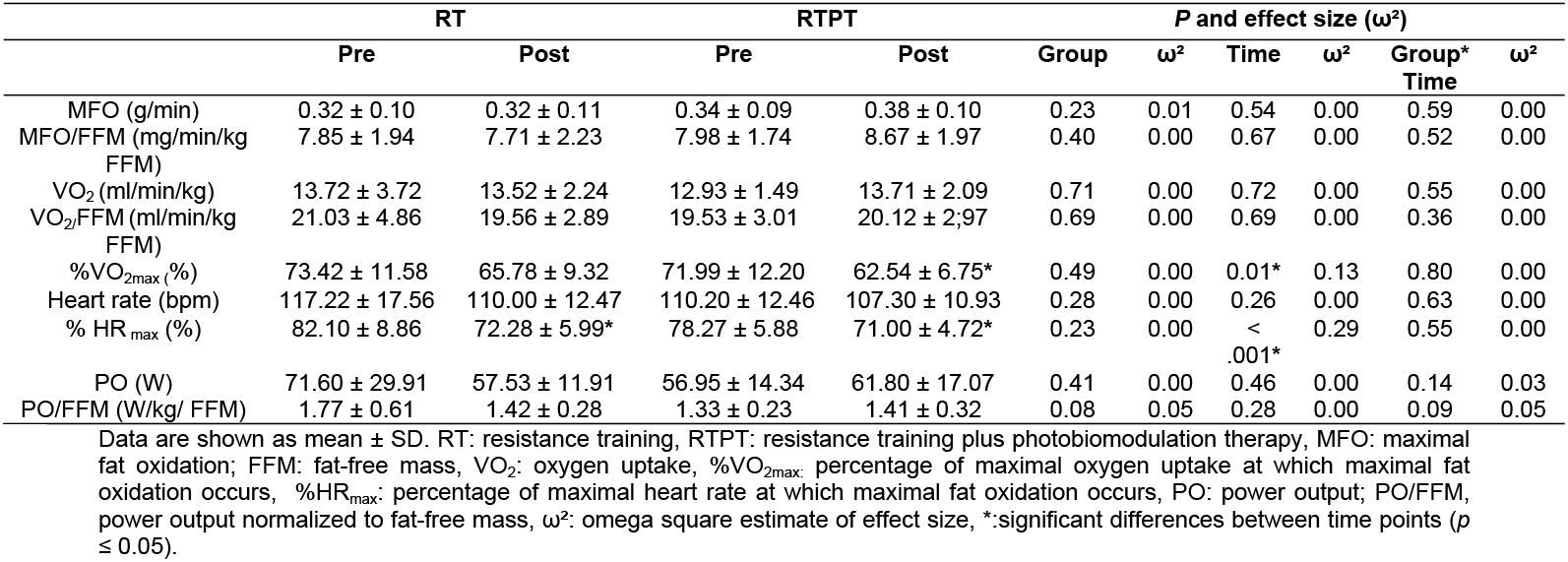
Comparison between groups in pre and post intervention at maximal fat oxidation.

### Carbohydrate and fat oxidation normalized to fat-free mass

No interaction between group*time on S1 FATox/FFM (*F*(1, 34) = 0.005, *p* = 0.94), S2 FATox/FFM (*F*(1, 34) = 0.95, *p* = 0.34), S3 FATox/FFM (*F*(1, 34) = 0.64, *p* = 0.43), S1 CHox/FFM (*F*(1, 34) = 1.19, *p* = 0.28), S2 CHox/FFM (I(1, 34) = 0.21, *p* = 0.65), and S3 CHox/FFM (*F*(1, 34) = 0.98, *p* = 0.33). However, a main effect of group was verified for S1 FATox/FFM (*F*(1, 34) = 6.69, *p* = 0.01), a main effect of time for S3 FATox/FFM (*F*(1, 34) = 9.32, *p* = 0.004), a main effect of time for S2 CHox/FFM (*F*(1, 34) = 5.70, *p* = 0.02), and a main effect of time for S3 CHox/FFM (*F*(1, 34) = 10.29, *p* = 0.002) was observed (Table 4).

**Table 4.** FATox/FFM and Chox/FFM in the incremental exercise test in pre and post intervention.

| | RT | | RTPT | | Group | P and effect size ($\omega^2$) | | | | |
| --- | --- | --- | --- | --- | --- | --- | --- | --- | --- | --- |
| | Pre | Post | Pre | Post | | $\omega^2$ | Time | $\omega^2$ | Group* Time | $\omega^2$ |
| **FATox/FFM (mg/min/kg FFM)** |  |  |  |  |  |  |  |  |  |  |
| Stage 1 | 4.72 ± 2.76 | 4.92 ± 1.86 | 6.64 ± 2.16 | 6.95 ± 2.51 | 0.01‡ | 0.14 | 0.74 | 0.00 | 0.94 | 0.00 |
| Stage 2 | 3.84 ± 2.49 | 2.18 ± 1.30 | 2.68 ± 1.73 | 2.14 ± 1.38 | 0.30 | 0.00 | 0.07 | 0.06 | 0.34 | 0.00 |
| Stage 3 | 0.26 ± 4.01 | -2.69 ± 2.72 | 0.14 ± 3.14 | -4.90 ± 5.54* | 0.38 | 0.00 | 0.004* | 0.18 | 0.43 | 0.00 |
| **CHox/FFM (mg/min/kg FFM)** |  |  |  |  |  |  |  |  |  |  |
| Stage 1 | 4.34 ± 2.16 | 8.21 ± 4.58 | 6.08 ± 4.53 | 6.84 ± 5.39 | 0.90 | 0.00 | 0.11 | 0.04 | 0.28 | 0.00 |
| Stage 2 | 21.49 ± 11.62 | 29.60 ± 9.00 | 26.00 ± 6.96 | 31.49 ± 7.03 | 0.27 | 0.00 | 0.02‡ | 0.11 | 0.65 | 0.00 |
| Stage 3 | 36.07 ± 16.93 | 46.51 ± 11.10 | 36.12 ± 12.28 | 55.92 ± 16.74* | 0.32 | 0.00 | 0.002* | 0.20 | 0.33 | 0.00 |

Data are shown as mean ± SD. RT: resistance training, RTPT: resistance training plus photobiomodulation therapy. Stage 1, stage 2 and stage 3 correspond to anaerobic threshold, respiratory compensation point and maximal oxygen uptake, respectively. $\omega^2$ : omega square estimate of effect size. \*:significant differences between time points ( $p \leq 0.05$ ), ‡: statistical difference became null after pairwise comparisons with Bonferroni correction.

After pairwise comparison with Bonferroni correction, differences became null (*p* = 0.084) for RT, and RTPT (*p* = 0.069), but with a small effect size for S1 FATox/FFM and S2 CHox/FFM (*p* = 0.058) for RT. For S3 FATox/FFM the RTPT displayed a statistically inferior mean difference compared to baseline of -5.04 (*p* = 0.008). Also, for S3 CHox/FFM the RTPT displayed a statistically superior mean difference compared to baseline of 19.80 (*p* = 0.004, and small effect size) (Table 4).

### Energy expenditure normalized to fat-free mass

No interaction between group*time on S1 EE/FFM (Cal/min/FFM) (*F*(1, 34) = 1.19, *p* = 0.28), S2 EE/FFM (Cal/min/FFM) (*F*(1, 34) = 0.21, *p* = 0.65), and S3 EE/FFM (Cal/min/FFM) (*F*(1, 34) = 0.98, *p* = 0.33). However, a main effect of time was verified for S2 EE/FFM (Cal/min/FFM) (*F*(1, 34) = 5.69, *p* = 0.02), and S3 EE/FFM (*F*(1, 34) = 10.29, *p* = 0.004) (Table 5).

**Table 5.** EE/FFM during incremental exercise test in pre and post intervention.

| | RT | | RTPT | | Group | P and effect size ($\omega^2$) | | | | |
| --- | --- | --- | --- | --- | --- | --- | --- | --- | --- | --- |
| | Pre | Post | Pre | Post | | $\omega^2$ | Time | $\omega^2$ | Group* Time | $\omega^2$ |
| **EE/FFM (Cal/min/FFM)** |  |  |  |  |  |  |  |  |  |  |
| Stage 1 | 17.71 ± 8.77 | 33.47 ± 18.63 | 24.80 ± 18.45 | 27.90 ± 21.93 | 0.90 | 0.00 | 0.11 | 0.04 | 0.28 | 0.00 |
| Stage 2 | 87.51 ± 47.29 | 120.48 ± 36.62 | 105.86 ± 28.33 | 128.19 ± 28.61 | 0.27 | 0.00 | 0.02‡ | 0.11 | 0.65 | 0.00 |
| Stage 3 | 146.79 ± 68.79 | 189.26 ± 45.16 | 147.00 ± 49.95 | 227.56 ± 68.08* | 0.32 | 0.00 | 0.004* | 0.20 | 0.33 | 0.00 |

Data are shown as mean ± SD. RT: resistance training, RTPT: resistance training plus photobiomodulation therapy. Stage 1, stage 2 and stage 3 correspond to anaerobic threshold, respiratory compensation point and maximal oxygen uptake, respectively. $\omega^2$ : omega square estimate of effect size, significant differences between time points ( $p \leq 0.05$ ), ‡: statistical difference became null after pairwise comparisons with Bonferroni correction.

After pairwise comparison with Bonferroni correction, differences became null (*p* = 0.058), but with a small effect size for S2 EE/FFM (Cal/min/FFM). For S3 EE/FFM (Cal/min/FFM) the RTPT displayed a statistically superior mean difference compared to baseline of 80.55 (*p* = 0.004) (Table 5).

## Discussion

In the present study, we evaluated the metabolic flexibility of sedentary elderly women in response to RT plus PBMT and RT alone through an incremental exercise test before and after 24 sessions of intervention. The key findings of the present study were that, in comparison with baseline RTPT displayed significant difference in VO_2max_, CHox, CHox/FFM and EE/FFM during maximum effort incremental testing. Moreover, there was a reduction in %VO_2max_ and %HR_max_ during maximum fat oxidation (MFO) for the RTPT, while RT group showed a significant difference only in the reduction of %HR_max_ during MFO.

These findings suggest that RT plus PBMT seems to have some additional positive impact on metabolic variables in elderly women when compared with RT alone. This confirms that exercise can be an important preventive strategy to improve metabolic flexibility and promote healthy aging [8], especially whether associated with PBMT, that seems to be a viable alternative to improve muscular performance and reduce the recovery time between exercise sessions. These benefits were previously observed in untrained subjects and athletes [17,18]. The PBMT acts directly on muscle tissue through the absorption of light by chromophores in cells, being cytochrome c oxidase (Cox) [19,20,47]. Cox is a complex IV in the mitochondrial respiratory chain that reduces oxygen to water, being considered the regulatory center of oxidative phosphorylation [48]. Thus, in theory, light coming from PBMT could dissociate nitric oxide (NO), the inhibitor of Cox, and increasing the electron transport, uptake oxygen and even gene expressions and cell proliferation [24,49].

It is known that PBMT can increase mitochondrial activity, resulting in greater ATP production and consequently better utilization of energy substrates [23,50], such as CHox and FATox, in this way, activating AMPK in the long term [14,25]. AMPK has also been considered an important regulator of exercise-induced effects on metabolic flexibility [27,28]. Although our results showed a significant difference in maximum oxygen consumption, CHox, and FATox when intensity was increased in the incremental test, observed in the RTPT group after the intervention compared to the RT group, we cannot say for sure whether these observations are due to changes at the mitochondrial level, because there were no assessments at the molecular or biochemical level. However, we report these results for the first time, and a more mechanistic approach should be conducted in future studies.

Both resistance and endurance training can provide considerable metabolic health benefits, such as increased mitochondrial content [14]. Furthermore, exercise training can partially reverse the age-related physiological decline and enhance work capacity in the elderly [51]. This can be especially important in the elderly, who have impaired physiological function. Thus, this stimulus with exercise training seems to result in faster positive metabolic effects in this population.

We believe that our results reflect this scenario, since both groups, RT and RTPT displayed positive changes, even if subtle, in some metabolic variables, but with a significant difference in favor of RTPT, likely due to the intervention with PBMT, which, as mentioned previously, acts directly on the mitochondria, promoting metabolic adaptations typical of aerobic training, such as mitochondrial dynamics and regulation of oxidative stress, collectively improving muscle performance and metabolic health [24,52,53].

Thus, PBMT can be considered as a possible potentiator of training-induced responses, as demonstrated in previous studies with the elderly population, both in animal models [54] and in humans [55,56]. However, the results should be interpreted with caution, due to differences in the protocols used among the studies. PBMT and exercise appear to combine well, helping to improve muscle function and to control inflammation, especially in older populations. Based on this, one could raise a pertinent question: could an intervention with elderly women combining RT and aerobic training have similar results in metabolic flexibility as the combination of RT and PBMT used in the present study? Future studies will be needed to answer this question.

The responses in VO2_max,_ CHox (Table 2), CHox/FFM (Table 4) as compared with baseline values during maximum effort (VO_2max_) incremental testing (S3) in the RTPT group seem to reflect the already positive description of the physical training combination with PBMT on metabolic adaptations. Substrate oxidation and use of oxygen are directly related to metabolic function, while the better capacity to oxidize substrates, the better the physical performance. The body has mechanisms to maintain energy homeostasis by adjusting the energy demand caused by exercise intensity. Thus, subjects whose metabolism prioritizes FATox and maintain basal lactate concentration at low values, and begin to prioritize CHox and increase lactate concentrations at higher intensities (e.g. end of effort test) present greater metabolic flexibility [11]. These mixed results suggest that PBMT combined with physical training show promise for enhancing metabolic adaptations in the elderly, while further research is needed to fully understand its efficacy and optimal light parameters application, including also different devices, such as those able to irradiate the whole body [57] and stimulate as much muscle mass as possible before or after the training sessions, as suggested previously [38] as well as a standardization between the protocols used.

In addition, MFO was observed at a lower %VO_2max_ (-9.34 %VO_2max_) in the RTPT group and at a lower %HR_max_ in both the RT and RTPT groups (-9.82 and -7,26 %HR_max_, respectively) when compared with baseline values (Table 3), without significantly changed for MFO in relation to baseline values. Like this, we attribute this increase in VO_2max_ to the intervention with PBMT, given that the combination of exercise with PBMT is known for its adaptations in energy metabolism, such as increased ATP production and improved utilization of energy substrates [50}.

It has been suggested that MFO can be increased by both aerobic or interval training, especially in sedentary populations (our sample), probably due to the adaptations inherent to aerobic metabolism with regard to fat oxidation, which is highly dependent on the delivery of fatty acids to skeletal muscle or the uptake of mitochondrial fatty acids [58-60]. Thus, the RT used in the present study may not have been sufficient to produce the metabolic adaptations expected with the combination proposed, as PBMT effects depend on the light dose and the time after application [17,23]. Furthermore, it is worth highlighting that MFO can be determined by several factors, including: training status, sex, acute nutritional status, chronic nutritional status, and possibly by the effect of exercise modality [60].

Despite the lack of change in absolute MFO (0.32 ± 0.1 and 0.38 ± 0.1 g/min for RT and RTPT, respectively), previous data in recreationally-active lean females revealed a value of 0.35 ± 0.12 g/min [60]. Moreover, MFO/FFM (mg/min/kg FFM), values were 7.71 ± 2.23 and 8.67 ± 1.97 mg/min/kg FFM, for RT and RTPT, respectively) values above those found by Blasco-Lafarga et al. [61] in physically active elderly women after an incremental test on a cycle ergometer (6.35 ± 3.59 mg/min/kg FFM). Elderly women tend to metabolic inflexibility, with little substrate availability and slower exchange between FATox and CHox in response to increasing intensity during incremental exercise [61]. However, findings from the present study suggest that the elderly women studied have a reasonable MFO compared to their peers from other studies, while there was no significant change in this variable after the intervention.

There was a considerable increase in EE/FFM for RTPT in the maximal effort test (S3) as compared with baseline, corresponding to 54.8% increase in the EE. This is due to the greater CHox/FFM observed in S3 and the increase in both EE/FFM and CHox. This increase could be associated with PBMT, that can increase glycogen synthesis in muscle tissue, providing an available eminent source of energy for use during aerobic activity [62], as observed in the final stages of an incremental maximal effort test, which may have allowed the elderly women to have greater energy expenditure and even achieve higher PO levels (Table 2).

The comparison of PO values of elderly women with other studies revealed that they present values above their counterparts as evidenced previously [61,63]. According to Blasco-Lafarga et al.[61] it is not aging that determines the drop in MFO, but muscle power, which seems to indicate that women who preserve power achieve a greater test performance, preserving the ability to oxidize more fat at lower intensities. Finally, muscle power has been identified as an extremely important variable for life expectancy and quality of life in the elderly, thus becoming a determining factor in the health of older women [61,64].

Among the strengths of study, we can highlight: first, the study uses a single-blind randomized controlled design with a placebo (sham) group, which allowed for greater control in the development of the research. Second, objective and high-quality physiological measurements, such as the use direct gas analysis (VO_2_, VCO_2_), ECG monitoring, and an incremental treadmill test. allowing detailed assessment across exercise intensities (AT, RCP, VO_2max_). Third, innovative research topic addressing a clear gap, the study explores the combination of resistance training plus photobiomodulation therapy on metabolic flexibility, an area with very limited prior research, especially in elderly women. Finally, well-controlled and clearly described intervention. Both the resistance training and PBMT protocols are well detailed and standardized, including: training intensity progression (70–100% of 10RM) and precise PBMT parameters (wavelength, dose, duration)

Finally, we highlight some limitations of the present study. First, our investigation presents a small sample, while we believe it is compensated by homogeneity of the characteristics, which made the comparison between groups more reliable. However, caution is advised when interpreting and extrapolating the results found here. Second, the lack of basal indirect calorimetry measurements to know the oxidation of substrates under resting conditions, prevented us from comparing the metabolic responses at rest with those during exercise. Third, we lacked a controlled nutritional intervention, although we believe that the pre-test dietary instructions were sufficient to avoid the influence on the results and that the self-selected diets are sufficient and reflect real life. Fourth, the TPBM intervention was performed only on the quadriceps femoris, despite the training being for the full body, limiting the magnitude of the PBMT effects; however, the focus of the study was precisely this: to specifically verify the response of TPBM in muscles important for activities of daily living in elderly, such as the quadriceps, one of the muscle groups most demanded in the incremental test. To note, the experimental design was controlled in the proposed aspects and included an unprecedented study in the literature.

## Conclusions

The intervention with RT plus PBMT seems to result in a positive impact on metabolic variables in sedentary elderly women when compared with RT alone, making this approach a viable alternative to promote positive changes in VO_2amx_, CHox, CHox/FFM and EE/FFM during maximal effort in incremental testing, as well as to reduce %VO_2max_ and %HR_max_ in MFO after 24 sessions. The combination of exercise plus PBMT may be recommended to promote positive changes in metabolic flexibility in elderly women.

Elderly women demonstrated high PO values compared to their age group, reinforcing that they present reasonable metabolic flexibility, considering the relationship between PO and mitochondrial adaptive responses. Finally, future studies with a similar approach will be necessary to better understand the metabolic responses arising from the combination of exercises with PBMT.

## Supporting information

**S1 Fig. Overview of the study design**

(PNG)

## Acknowledgements

We express our sincere gratitude to PhD Dahan da Cunha Nascimento for their valuable insights regarding the statistical analysis.

## Author Contributions

**Conceptualization:** Ronaldo Ferreira Moura, Emanuelle Fernandes Prestes, Danielle Garcia de Araujo, Jonato Prestes.

**Data curation:** Ronaldo Ferreira Moura, Emanuelle Fernandes Prestes, Danielle Garcia de Araujo, Jonato Prestes.

**Investigation:** Gislane Ferreira de Melo, Thiago dos Santos Rosa, James W. Navalta, Wilson Max Almeida Monteiro de Moraes, Cleber Ferraresi.

**Methodology:** Ronaldo Ferreira Moura, Emanuelle Fernandes Prestes, Danielle Garcia de Araujo, Jonato Prestes.

**Supervision:** James W. Navalta, Cleber Ferraresi.

**Writing – original draft:** Ronaldo Ferreira Moura, Emanuelle Fernandes Prestes, Danielle Garcia de Araujo.

**Writing – review & editing:** Gislane Ferreira de Melo, Thiago dos Santos Rosa, Wilson Max Almeida Monteiro de Moraes, Joanto Prestes

## References

1. Goodpaster BH, Sparks LM. Metabolic Flexibility in Health and Disease. Cell Metab. 2017 May 2;25(5):1027–1036. doi: 10.1016/j.cmet.2017.04.015. PMID: 28467922.

2. Smith RL, Soeters MR, Wüst RCI, Houtkooper RH. Metabolic Flexibility as an Adaptation to Energy Resources and Requirements in Health and Disease. Endocr Rev. 2018 Aug 1;39(4):489–517. doi: 10.1210/er.2017-00211. PMID: 29697773.

3. Benítez-Muñoz JA, Guisado-Cuadrado I, Rojo-Tirado M, Alcocer-Ayuga M, Romero-Parra N, Peinado AB, Cupeiro R. Females have better metabolic flexibility in different metabolically challenging stimuli. Appl Physiol Nutr Metab. 2025 Jan 1;50:1–12. doi: 10.1139/apnm-2024-0217. PMID: 39437435.

4. Garthwaite T, Sjöros T, Laine S, Koivumäki M, Vähä-Ypyä H, Verho T, Norha J, Kallio P, Saarenhovi M, Löyttyniemi E, Sievänen H, Houttu N, Laitinen K, Kalliokoski KK, Vasankari T, Knuuti J, Heinonen I. Sedentary time associates detrimentally and physical activity beneficially with metabolic flexibility in adults with metabolic syndrome. Am J Physiol Endocrinol Metab. 2024 Apr 1;326(4):E503–E514. doi:10.1152/ajpendo.00338.2023. PMID: 38416072.

5. Shoemaker ME, Pereira SL, Mustad VA, Gillen ZM, McKay BD, Lopez-Pedrosa JM, Rueda R, Cramer JT. Differences in muscle energy metabolism and metabolic flexibility between sarcopenic and nonsarcopenic older adults. J Cachexia Sarcopenia Muscle. 2022 Apr;13(2):1224–1237. doi: 10.1002/jcsm.12932. PMID: 35178889.

6. Wiley CD, Campisi J. From Ancient Pathways to Aging Cells-Connecting Metabolism and Cellular Senescence. Cell Metab. 2016 Jun 14;23(6):1013–1021. doi: 10.1016/j.cmet.2016.05.010. PMID: 27304503.

7. Wiley CD, Velarde MC, Lecot P, Liu S, Sarnoski EA, Freund A, Shirakawa K, Lim HW, Davis SS, Ramanathan A, Gerencser AA, Verdin E, Campisi J. Mitochondrial Dysfunction Induces Senescence with a Distinct Secretory Phenotype. Cell Metab. 2016 Feb 9;23(2):303–14. doi: 10.1016/j.cmet.2015.11.011. PMID: 26686024.

8. Cartee GD, Hepple RT, Bamman MM, Zierath JR. Exercise Promotes Healthy Aging of Skeletal Muscle. Cell Metab. 2016 Jun 14;23(6):1034–1047. doi: 10.1016/j.cmet.2016.05.007. PMID: 27304505.

9. Battaglia GM, Zheng D, Hickner RC, Houmard JA. Effect of exercise training on metabolic flexibility in response to a high-fat diet in obese individuals. Am J Physiol Endocrinol Metab. 2012 Dec 15;303(12):E1440–5. doi: 10.1152/ajpendo.00355.2012. PMID: 23047988.

10. Amaro-Gahete FJ, Sanchez-Delgado G, Jurado-Fasoli L, De-la-O A, Castillo MJ, Helge JW, Ruiz JR. Assessment of maximal fat oxidation during exercise: A systematic review. Scand J Med Sci Sports. 2019 Jul;29(7):910–921. doi: 10.1111/sms.13424. PMID: 30929281.

11. San-Millán I, Brooks GA. Assessment of Metabolic Flexibility by Means of Measuring Blood Lactate, Fat, and Carbohydrate Oxidation Responses to Exercise in Professional Endurance Athletes and Less-Fit Individuals. Sports Med. 2018 Feb;48(2):467–479. doi: 10.1007/s40279-017-0751-x. PMID: 28623613.

12. Fernández-Verdejo R, Gutiérrez-Pino J, Hayes-Ortiz T, Zbinden-Foncea H, Cabello-Verrugio C, Valero-Breton M, Tuñón-Suárez M, Vargas-Foitzick R, Galgani JE. Metabolic flexibility to lipid during exercise is not associated with metabolic health outcomes in individuals without obesity. Sci Rep. 2024 Nov 19;14(1):28642. doi: 10.1038/s41598-024-79092-w. PMID: 39562632.

13. Ortmeyer HK, Goldberg AP, Ryan AS. Exercise with weight loss improves adipose tissue and skeletal muscle markers of fatty acid metabolism in postmenopausal women. Obesity (Silver Spring). 2017 Jul;25(7):1246–1253. doi: 10.1002/oby.21877. PMID: 28547918.

14. Egan B, Zierath JR. Exercise metabolism and the molecular regulation of skeletal muscle adaptation. Cell Metab. 2013 Feb 5;17(2):162–84. doi: 10.1016/j.cmet.2012.12.012. PMID: 23395166.

15. Sene-Fiorese M, Duarte FO, de Aquino Junior AE, Campos RM, Masquio DC, Tock L, de Oliveira Duarte AC, Dâmaso AR, Parizotto NA, Bagnato VS. The potential of phototherapy to reduce body fat, insulin resistance and “metabolic inflexibility” related to obesity in women undergoing weight loss treatment. Lasers Surg Med. 2015 Oct;47(8):634–42. doi: 10.1002/lsm.22395. PMID: 26220050.

16. Ferraresi C, Hamblin MR, Parizotto NA. Low-level laser (light) therapy (LLLT) on muscle tissue: performance, fatigue and repair benefited by the power of light. Photonics Lasers Med. 2012 Nov 1;1(4):267–286. doi: 10.1515/plm-2012-0032. PMID: 23626925.

17. Ferraresi C, Huang YY, Hamblin MR. Photobiomodulation in human muscle tissue: an advantage in sports performance? J Biophotonics. 2016 Dec;9(11-12):1273–1299. doi: 10.1002/jbio.201600176. PMID: 27874264.

18. Vanin AA, Verhagen E, Barboza SD, Costa LOP, Leal-Junior ECP. Photobiomodulation therapy for the improvement of muscular performance and reduction of muscular fatigue associated with exercise in healthy people: a systematic review and meta-analysis. Lasers Med Sci. 2018 Jan;33(1):181–214. doi: 10.1007/s10103-017-2368-6. PMID: 29090398.

19. Karu T. Primary and secondary mechanisms of action of visible to near-IR radiation on cells. J Photochem Photobiol B. 1999 Mar;49(1):1–17. doi: 10.1016/S1011-1344(98)00219-X. PMID: 10365442.

20. Karu TI, Pyatibrat LV, Kolyakov SF, Afanasyeva NI. Absorption measurements of a cell monolayer relevant to phototherapy: reduction of cytochrome c oxidase under near IR radiation. J Photochem Photobiol B. 2005 Nov 1;81(2):98–106. doi: 10.1016/j.jphotobiol.2005.07.002. PMID: 16125966.

21. Anders JJ, Lanzafame RJ, Arany PR. Low-level light/laser therapy versus photobiomodulation therapy. Photomed Laser Surg. 2015 Apr;33(4):183–4. doi: 10.1089/pho.2015.9848. PMID: 25844681.

22. Heiskanen V, Hamblin MR. Photobiomodulation: lasers vs. light emitting diodes? Photochem Photobiol Sci. 2018 Aug 8;17(8):1003–1017. doi: 10.1039/c8pp90049c. Erratum in: Photochem Photobiol Sci. 2018 Oct 31;18(1):259-259. doi: 10.1039/c8pp90049c. PMID: 30044464.

23. Ferraresi C, Kaippert B, Avci P, Huang YY, de Sousa MV, Bagnato VS, Parizotto NA, Hamblin MR. Low-level laser (light) therapy increases mitochondrial membrane potential and ATP synthesis in C2C12 myotubes with a peak response at 3-6 h. Photochem Photobiol. 2015 Mar-Apr;91(2):411–6. doi: 10.1111/php.12397. PMID: 25443662.

24. Hamblin MR. Mechanisms and Mitochondrial Redox Signaling in Photobiomodulation. Photochem Photobiol. 2018 Mar;94(2):199–212. doi: 10.1111/php.12864. PMID: 29164625.

25. Guo S, Gong L, Shen Q, Xing D. Photobiomodulation reduces hepatic lipogenesis and enhances insulin sensitivity through activation of CaMKKβ/AMPK signaling pathway. J Photochem Photobiol B. 2020 Dec;213:112075. doi: 10.1016/j.jphotobiol.2020.112075. PMID: 33152638.

26. Ke R, Xu Q, Li C, Luo L, Huang D. Mechanisms of AMPK in the maintenance of ATP balance during energy metabolism. Cell Biol Int. 2018 Apr;42(4):384–392. doi: 10.1002/cbin.10915. PMID: 29205673.

27. Kjøbsted R, Munk-Hansen N, Birk JB, Foretz M, Viollet B, Björnholm M, Zierath JR, Treebak JT, Wojtaszewski JF. Enhanced Muscle Insulin Sensitivity After Contraction/Exercise Is Mediated by AMPK. Diabetes. 2017 Mar;66(3):598–612. doi: 10.2337/db16-0530. PMID: 27797909.

28. Mambrini SP, Grillo A, Colosimo S, Zarpellon F, Pozzi G, Furlan D, Amodeo G, Bertoli S. Diet and physical exercise as key players to tackle MASLD through improvement of insulin resistance and metabolic flexibility. Front Nutr. 2024 Aug 20;11:1426551. doi: 10.3389/fnut.2024.1426551. PMID: 39229589.

29. Bull FC, Al-Ansari SS, Biddle S, Borodulin K, Buman MP, Cardon G, Carty C, Chaput JP, Chastin S, Chou R, Dempsey PC, DiPietro L, Ekelund U, Firth J, Friedenreich CM, Garcia L, Gichu M, Jago R, Katzmarzyk PT, Lambert E, Leitzmann M, Milton K, Ortega FB, Ranasinghe C, Stamatakis E, Tiedemann A, Troiano RP, van der Ploeg HP, Wari V, Willumsen JF. World Health Organization 2020 guidelines on physical activity and sedentary behaviour. Br J Sports Med. 2020 Dec;54(24):1451–1462. doi: 10.1136/bjsports-2020-102955. PMID: 33239350.

30. World Medical Association. World Medical Association Declaration of Helsinki: ethical principles for medical research involving human subjects. JAMA. 2013 Nov 27;310(20):2191–4. doi: 10.1001/jama.2013.281053. PMID: 24141714.

31. Sharman JE, LaGerche A. Exercise blood pressure: clinical relevance and correct measurement. J Hum Hypertens. 2015 Jun;29(6):351–8. doi: 10.1038/jhh.2014.84. PMID: 25273859.

32. Wasserman K, Hansen JE, Sue DY, Stringer WW, Whipp BJ. Principles of exercise testing and interpretation: including pathophysiology and clinical applications. Med. Sci. Sports Exerc. 2005 jul;37(7); 1249. doi: 10.1249/01.mss.0000172593.20181.14.

33. Beaver WL, Wasserman K, Whipp BJ. A new method for detecting anaerobic threshold by gas exchange. J Appl Physiol (1985). 1986 Jun;60(6):2020–7. doi: 10.1152/jappl.1986.60.6.2020. PMID: 3087938.

34. Kominami K, Imahashi K, Katsuragawa T, Murakami M, Akino M. The Ratio of Oxygen Uptake From Ventilatory Anaerobic Threshold to Respiratory Compensation Point Is Maintained During Incremental Exercise in Older Adults. Front Physiol. 2022 Mar 3;13:769387. doi: 10.3389/fphys.2022.769387. PMID: 35309068.

35. Poole DC, Wilkerson DP, Jones AM. Validity of criteria for establishing maximal O2 uptake during ramp exercise tests. Eur J Appl Physiol. 2008 Mar;102(4):403–10. doi: 10.1007/s00421-007-0596-3. PMID: 17968581.

36. Robinson MM, Dasari S, Konopka AR, Johnson ML, Manjunatha S, Esponda RR, Carter RE, Lanza IR, Nair KS. Enhanced Protein Translation Underlies Improved Metabolic and Physical Adaptations to Different Exercise Training Modes in Young and Old Humans. Cell Metab. 2017 Mar 7;25(3):581–592. doi: 10.1016/j.cmet.2017.02.009. PMID: 28273480.

37. Neto RPM, Espósito LMB, da Rocha FC, Filho AAS, Silva JHG, de Sousa Santos EC, Sousa BLSC, Dos Santos Gonçalves KRR, Garcia-Araujo AS, Hamblin MR, Ferraresi C. Photobiomodulation therapy (red/NIR LEDs) reduced the length of stay in intensive care unit and improved muscle function: A randomized, triple-blind, and sham-controlled trial. J Biophotonics. 2024 May;17(5):e202300501. doi: 10.1002/jbio.202300501. PMID: 38262071.

38. Ferraresi C. Use of Photobiomodulation Therapy in Exercise Performance Enhancement and Postexercise Recovery: True or Myth? Photobiomodul Photomed Laser Surg. 2020 Dec;38(12):705–707. doi: 10.1089/photob.2020.4948. PMID: 33216681.

39. Ferraresi C, Beltrame T, Fabrizzi F, do Nascimento ES, Karsten M, Francisco Cde O, Borghi-Silva A, Catai AM, Cardoso DR, Ferreira AG, Hamblin MR, Bagnato VS, Parizotto NA. Muscular pre-conditioning using light-emitting diode therapy (LEDT) for high-intensity exercise: a randomized double-blind placebo-controlled trial with a single elite runner. Physiother Theory Pract. 2015 Jul;31(5):354–61. doi: 10.3109/09593985.2014.1003118. PMID: 25585514.

40. Zamani ARN, Saberianpour S, Geranmayeh MH, Bani F, Haghighi L, Rahbarghazi R. Modulatory effect of photobiomodulation on stem cell epigenetic memory: a highlight on differentiation capacity. Lasers Med Sci. 2020 Mar;35(2):299–306. doi: 10.1007/s10103-019-02873-7. PMID: 31494789.

41. Frayn KN. Calculation of substrate oxidation rates in vivo from gaseous exchange. J Appl Physiol Respir Environ Exerc Physiol. 1983 Aug;55(2):628–34. doi: 10.1152/jappl.1983.55.2.628. PMID: 6618956.

42. Jeukendrup AE, Wallis GA. Measurement of substrate oxidation during exercise by means of gas exchange measurements. Int J Sports Med. 2005 Feb;26 Suppl 1:S28–37. doi: 10.1055/s-2004-830512. PMID: 15702454.

43. Fey CF, Hu T, Delios A. The measurement and communication of effect sizes in management research. Manage. Organ. Rev. 2023 Apr;19(1), 176–197.

44. Mayr S, Erdfelder E, Buchner A, Faul F. A short tutorial of GPower. Tutor in quant method for psychol. 2007;3(2): 51–9.

45. Beck TW. The importance of a priori sample size estimation in strength and conditioning research. J Strength Cond Res. 2013 Aug;27(8):2323–37. doi: 10.1519/JSC.0b013e318278eea0. PMID: 23880657.

46. Goss-Sampson M. Statistical analysis in JASP: A guide for students. JASP. 2019.

47. de Freitas LF, Hamblin MR. Proposed Mechanisms of Photobiomodulation or Low-Level Light Therapy. IEEE J Sel Top Quantum Electron. 2016 May-Jun;22(3):7000417. doi: 10.1109/JSTQE.2016.2561201. PMID: 28070154.

48. Kadenbach B. Complex IV - The regulatory center of mitochondrial oxidative phosphorylation. Mitochondrion. 2021 May;58:296–302. doi: 10.1016/j.mito.2020.10.004. PMID: 33069909.

49. Glass GE. Photobiomodulation: A review of the molecular evidence for low level light therapy. J Plast Reconstr Aesthet Surg. 2021 May;74(5):1050–1060. doi: 10.1016/j.bjps.2020.12.059. PMID: 33436333.

50. Hamblin MR. Mechanisms and applications of the anti-inflammatory effects of photobiomodulation. AIMS Biophys. 2017;4(3):337–361. doi: 10.3934/biophy.2017.3.337. PMID: 28748217.

51. Mendonca GV, Pezarat-Correia P, Vaz JR, Silva L, Almeida ID, Heffernan KS. Impact of Exercise Training on Physiological Measures of Physical Fitness in the Elderly. Curr Aging Sci. 2016;9(4):240–259. doi: 10.2174/1874609809666160426120600. PMID: 27113585.

52. Philp AM, Saner NJ, Lazarou M, Ganley IG, Philp A. The influence of aerobic exercise on mitochondrial quality control in skeletal muscle. J Physiol. 2021 Jul;599(14):3463–3476. doi: 10.1113/JP279411. PMID: 33369731.

53. Trajano LADSN, Siqueira PB, Rodrigues MMS, Pires BRB, da Fonseca AS, Mencalha AL. Does photobiomodulation alter mitochondrial dynamics? Photochem Photobiol. 2025 Jan-Feb;101(1):21–37. doi: 10.1111/php.13963. PMID: 38774941.

54. Guaraldo SA, Serra AJ, Amadio EM, Antônio EL, Silva F, Portes LA, Tucci PJ, Leal-Junior EC, de Carvalho Pde T. The effect of low-level laser therapy on oxidative stress and functional fitness in aged rats subjected to swimming: an aerobic exercise. Lasers Med Sci. 2016 Jul;31(5):833–40. doi: 10.1007/s10103-016-1882-2. PMID: 26861983.

55. Vassão PG, de Souza ACF, da Silveira Campos RM, Garcia LA, Tucci HT, Renno ACM. Effects of photobiomodulation and a physical exercise program on the expression of inflammatory and cartilage degradation biomarkers and functional capacity in women with knee osteoarthritis: a randomized blinded study. Adv Rheumatol. 2021 Oct 16;61(1):62. doi: 10.1186/s42358-021-00220-5. PMID: 34656170.

56. Kumar P, Umakanth S N G. Photobiomodulation therapy as an adjunct to resistance exercises on muscle metrics, functional balance, functional capacity, and physical performance among older adults: A systematic scoping review. Lasers Med Sci. 2024 Sep 3;39(1):232. doi: 10.1007/s10103-024-04177-x. PMID: 39225877.

57. Zagatto AM, Dutra YM, Lira FS, Antunes BM, Faustini JB, Malta ES, Lopes VHF, de Poli RAB, Brisola GMP, Dos Santos GV, Rodrigues FM, Ferraresi C. Full Body Photobiomodulation Therapy to Induce Faster Muscle Recovery in Water Polo Athletes: Preliminary Results. Photobiomodul Photomed Laser Surg. 2020 Dec;38(12):766–772. doi: 10.1089/photob.2020.4803. PMID: 33332232.

58. van Loon LJ, Greenhaff PL, Constantin-Teodosiu D, Saris WH, Wagenmakers AJ. The effects of increasing exercise intensity on muscle fuel utilisation in humans. J Physiol. 2001 Oct 1;536(Pt 1):295–304. doi: 10.1111/j.1469-7793.2001.00295.x. PMID: 11579177.

59. Spriet LL. New insights into the interaction of carbohydrate and fat metabolism during exercise. Sports Med. 2014 May;44 Suppl 1(Suppl 1):S87–96. doi: 10.1007/s40279-014-0154-1. PMID: 24791920.

60. Maunder E, Plews DJ, Kilding AE. Contextualising Maximal Fat Oxidation During Exercise: Determinants and Normative Values. Front Physiol. 2018 May 23;9:599. doi: 10.3389/fphys.2018.00599. PMID: 29875697.

61. Blasco-Lafarga C, Monferrer-Marín J, Roldán A, Monteagudo P, Chulvi-Medrano I. Metabolic Flexibility and Mechanical Efficiency in Women Over-60. Front Physiol. 2022 Apr 6;13:869534. doi: 10.3389/fphys.2022.869534. PMID: 35464093.

62. Castro KMR, de Paiva Carvalho RL, Junior GMR, Tavares BA, Simionato LH, Bortoluci CHF, Soto CAT, Ferraresi C. Can photobiomodulation therapy (PBMT) control blood glucose levels and alter muscle glycogen synthesis? J Photochem Photobiol B. 2020 Jun;207:111877. doi: 10.1016/j.jphotobiol.2020.111877. PMID: 32298941.

63. Monferrer-Marín J, Roldán A, Monteagudo P, Chulvi-Medrano I, Blasco-Lafarga C. Impact of Ageing on Female Metabolic Flexibility: A Cross-Sectional Pilot Study in over-60 Active Women. Sports Med Open. 2022 Jul 30;8(1):97. doi: 10.1186/s40798-022-00487-y. PMID: 35907092.

64. Zambom-Ferraresi F, Cebollero P, Gorostiaga EM, Hernández M, Hueto J, Cascante J, Rezusta L, Val L, Anton MM. Effects of Combined Resistance and Endurance Training Versus Resistance Training Alone on Strength, Exercise Capacity, and Quality of Life in Patients With COPD. J Cardiopulm Rehabil Prev. 2015 Nov-Dec;35(6):446–53. doi: 10.1097/HCR.0000000000000132. PMID: 26252342.

